# Structure of ER chaperone complex GRP170-ATP-BiP suggests a new model for substrate engagement

**DOI:** 10.1101/2025.10.25.684324

**Authors:** Yassin Ben-Khoud, Kono Washio, Srinjan Basu, Maruf MU Ali

## Abstract

Molecular chaperones are essential for maintaining protein homeostasis in all living cells^1^. In the endoplasmic reticulum (ER), BiP and GRP170 are the sole representatives of Hsp70 and Hsp110 family and are critical for ER function. GRP170 is a relatively large and unusual chaperone that possess both nucleotide exchange and chaperoning activity^2^. The molecular mechanism by which these chaperones collaborate to engage substrate protein and how GRP170 couples its dual functionalities are not currently known. Here, we report the 2.7 Å cryo-electron microscopy structure of GRP170-ATP-BiP chaperone complex purified from HEK293 cells that reveals a C-terminal curved hook domain, suggesting a role in substrate engagement in coordination with BiP. Additionally, we uncover the structural basis for GRP170 pseudo-ATPase chaperone activity – making it, to our knowledge, the first chaperone with this type of regulation. Our ER chaperone complex structure, together with prior cellular data^3^, suggests a new paradigm for how GRP170-BiP chaperones collaborate in ER protein quality control, broadening our understanding of how BiP/Hsp70 chaperones engage with substrates.

## Main

The endoplasmic reticulum is second only to the cytosol in the folding and maturation of proteins within the eukaryotic cell. As the first compartment in the secretory pathway, the ER plays a pivotal role in processing proteins destined for locations beyond the ER, including the extracellular space. Consequently, it is especially significant in the production and modification of secretory proteins, such as hormones and antibodies^4,5^.

Molecular chaperones are critical for ensuring nascent proteins attain their correct native shape and for preserving their structural integrity during their functional life span. There are several different families of chaperones and multiple cochaperones that combine to form chaperone complexes within each compartment of the cell. These complexes are vital for mediating the association with substrate proteins and ensuring successful chaperone function and cellular fitness^1^.

There are notable differences between the molecular chaperone systems of the cytosol and the ER, reflecting the distinct biochemical environment of these compartments. While the cytosol contains multiple isoforms of major chaperone families – such as Hsp70, Hsp90, and Hsp110 – the ER harbours only a single isoform of each. Furthermore, cochaperones that facilitate interactions between different chaperone families in the cytosol appear to be absent in the ER, suggesting that ER chaperones may operate via different mechanisms of action that remain incompletely understood^1,2^.

The sole ER Hsp70 chaperone, BiP, forms chaperone complexes that are fundamental to protein homeostasis within this organelle. Notably, BiP serves a much greater diversity of roles than its cytosolic counterparts, this includes protein translocation across the ER membrane, ER associated degradation (ERAD), UPR activation, as well as its canonical function in protein folding. BiP consists of a nucleotide binding domain (NBD) that hydrolyses ATP and a substrate binding domain (SBD) that engages misfolded protein substrates. BiP associates with two groups of cochaperones: 1) J-domain proteins that stimulate ATP hydrolysis and 2) Nucleotide exchange factors (NEF) that facilitate exchange of ADP to ATP^2,6,7^.

BiP interacts with a large and unusual ER-resident chaperone known as glucose regulated protein 170 (GRP170). GRP170 belongs to the Hsp110 family of chaperones, which are modelled on the cytosolic yeast homolog Sse1. Members of the Hsp110 family share structural similarities with Hsp70, particularly the NBD, but differ markedly in other regions. They feature an acidic insertion within the β-sandwich domain and an extended C-terminal α-helical domain^8^. Among its family, GRP170 is especially distinctive. Compared to cytosolic isoforms, GRP170 possesses a larger C-terminal extension and is extensively glycosylated^9^.

GRP170 is upregulated in response to various cytotoxic and proteotoxic stresses, including heat shock, viral infections, and inflammation^10^. Recent studies have linked GRP170 dysfunction to impaired ion homeostasis^11^. Notably, GRP170 ablation disrupts BiP function, triggers apoptosis, and causes severe loss of ER morphology^12^. These findings highlight BiP and GRP170 as the only ER chaperones essential for cell survival, underscoring their critical roles in maintaining cellular health and homeostasis.

GRP170 functions both as a bona fide molecular chaperone and as a NEF. As a chaperone, it binds misfolded proteins and protein assemblies to prevent their aggregation^10,11,13^. Unusually, GRP170’s interaction with client substrate is not disrupted by ATP—which is unlike any known Hsp70 mechanism^3^.

Recent structures of cytosolic complexes have generated mechanistic insights into Hsp70 and Hsp90 chaperone families^14–16^. Typically, they have been reconstituted and fixed with crosslinker to aid sample stabilization. While crystal structures of cytosolic Sse1 bound to Hsp70 NBD provide information of NEF interactions in yeast^17,18^, there is a lack of high-resolution structural data for ER-derived complexes from any of the chaperone families. Moreover, the structure and functional role of GRP170 C-terminal extension also remains unresolved, as does the mechanism by which it integrates its dual roles as a NEF and a molecular chaperone.

To advance our understanding of the molecular mechanisms underlying ER chaperone complexes, we purified GRP170-ATP-BiP complex directly from HEK293 cells and determined its structure by cryo-electron microscopy (cryo-EM). The resulting structure provides novel insights into the architecture and regulation of the chaperone complex, helping to clarify previously unexplained biochemical and cellular observations^3^. Collectively, these findings suggest a new paradigm for GRP170-BiP molecular function in ER homeostasis.

### Structure determination of ER chaperone complex

A construct encoding full-length GRP170 with a triple FLAG tag inserted between the native signal sequence and the mature protein was cloned into pcDNA3.1+ vector for expression in HEK293 suspension cells. During the purification, an ATP incubation step was included prior to elution with FLAG peptide. SDS-PAGE analysis revealed a prominent band corresponding to GRP170 (Extended Data Fig. 1), which was confirmed by mass spectrometry. A second less intense band at ∼70 kDa size was subsequently identified as endogenous BiP, indicating the purification of an ER chaperone complex directly from HEK293 cells in the presence of ATP.

Cryo-EM grids were prepared by vitrifying the protein sample in vitreous ice for single particle analysis. Initial 2D class averages revealed a stable particle that suggested the presence of two molecules. However, the lack of symmetry between the two components indicated that they were distinct proteins and not a dimer of GRP170. Subsequent cryo-EM reconstruction resolved the structure at a global resolution of 2.7 Å, clearly identifying density corresponding to endogenous BiP bound to GRP170.

### Structure of GRP170-ATP-BiP reveals hook domain

GRP170 and BiP form a 1:1 heterodimeric protein complex (Fig. 1). For GRP170, continuous density enabled atomic modelling for most of the protein including the NBD and β-sandwich domain, with exception of the acidic loop. The previously uncharacterised C-terminal α-helical domain was reasonably well resolved for the most part (minus the end ∼84 residues) after we performed a focused 3D classification and focused refinement to enhance signal and improve resolution (Extended Data Fig. 2, 3).

**Fig. 1:**
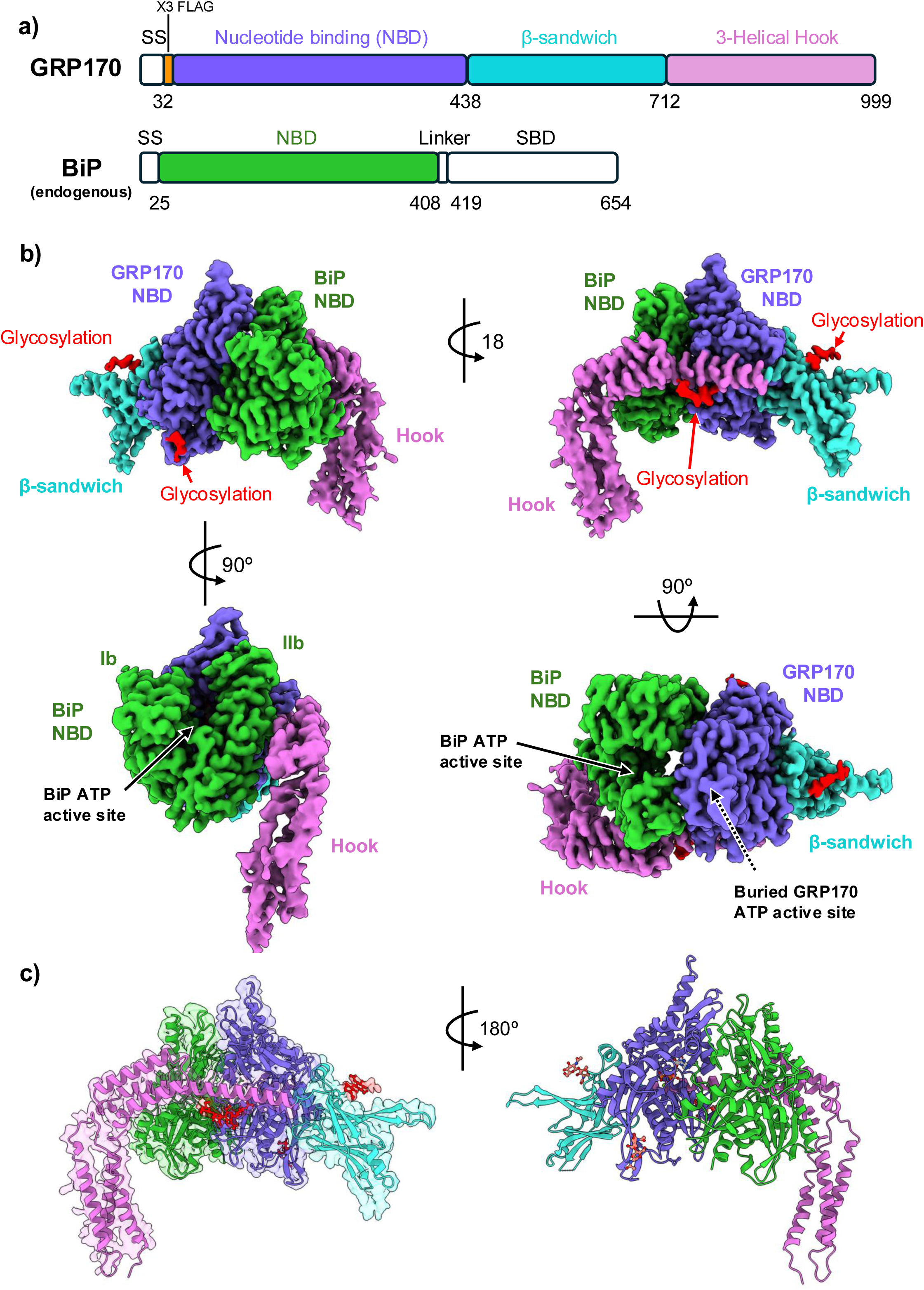
Structure of GRP170-ATP-BiP chaperone complex. **a**, domain organisation of GRP170 and BiP chaperones, with GRP170 NBD in purple, β-sandwich in cyan, hook domain in magenta, BiP NBD is green. **b**, Composite cryo-EM map of chaperone complex viewed from side, top and front view. The colours reflect the domains in (**a**), with glycan coloured in red. GRP170 C-terminal adopts a striking hook shape that is supported by contacts with BiP NBD and a glycan. The BiP NBD lobes Ib and IIb are labelled and show that the active site is relatively open and accessible to solvent, whilst GRP170 active site is buried and inaccessible, as indicated by dashed arrow. **c**, Atomic structure modelled into cryo-EM map.

The most striking feature of the complex structure is the way the α-helical domain forms a hook shape that folds back towards the GRP170 NBD (Fig. 1). The architecture of this hook domain is unusual and not observed in other chaperone families including Hsp70 and Hsp90. Earlier work has shown that this domain was able to bind to misfolded substrate in an ATP independent fashion^3^, though it was unclear how this would occur. Considering this prior important observation, the curved hook domain architecture suggests that it may collaborate with BiP SBD to engage substrate protein.

GRP170 NBD consists of 4 lobes that are arranged to form a central cavity for nucleotide binding and a β-sandwich domain comprising two-layer anti-parallel β-sheets that has a similar fold to NBD and SBD from both Hsp110 and Hsp70, as expected. Similarly, BiP NBD forms the classical fold of Hsp70 NBD. Putative density corresponding to BiP SBD was only visible following 3D variability analysis (3DVA) focused on BiP SBD and further enhanced by contouring at low map threshold. However, the resulting density was not of sufficient quality to model (Extended Data Fig. 3H). Nonetheless, it suggests a close connection between the hook domain and BiP SBD. There are three N-glycans that have been modelled, with one that supports the hook domain extension (Extended Data Fig. 4).

An overview of the heterodimer structure indicates that the BiP NBD is relatively open, and access to the ATPase active site is unrestricted, whilst GRP170 active site is quite buried, and formation of the complex would restrict any access to the active site (Fig. 1). Structurally, to exchange nucleotide from the GRP170 active site would require breakdown of the heterodimer complex, while exchange of nucleotide from Hsp70 can occur whilst bound to GRP170.

### The large heterodimer interface is conserved

The interaction between BiP and GRP170 involves a substantial interface, burying approximately 3150 Å² of total surface area. This interface comprises two distinct regions, resembling the Sse1-Hsp70 interaction^17,18^, with each region contributing to the stability and specificity of the heterodimer complex (Fig. 2).

**Fig. 2:**
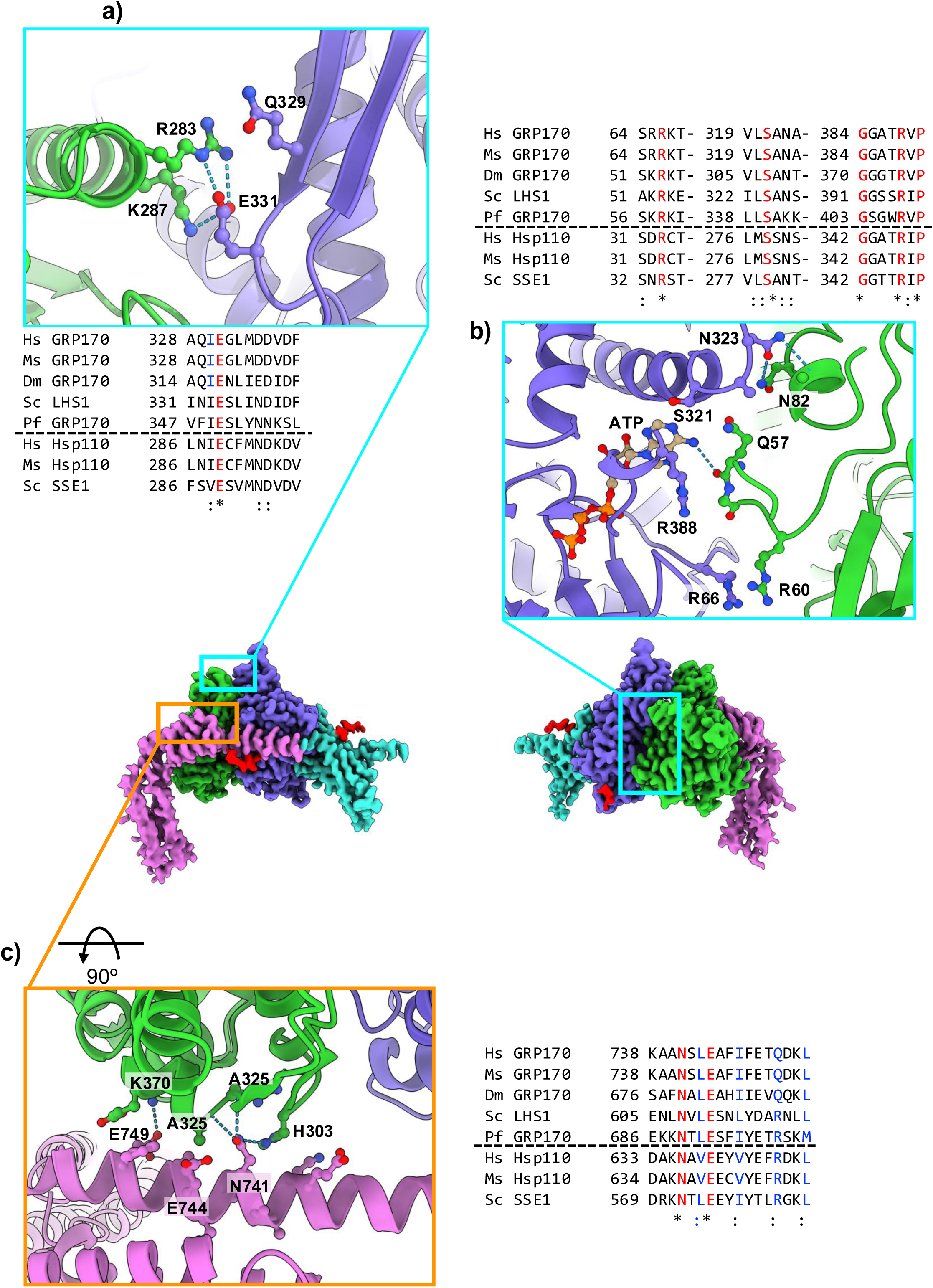
Molecular interactions between chaperones in the complex. There are two distinct interface regions -Interface 1: a face-to-face interaction between NBD of GRP170 and BiP (frame coloured in cyan). Interface 2: BiP NBD association with the hook domain (frame coloured in orange). **a**, Illustrates the upper region of interface 1, highlighting hydrogen bonds formed between the conserved E331 of GRP170 and BiP NBD. **b**, shows the core interactions of interface 1 comprising substantial van der Waals contacts and several hydrogen bonds. Key residues include the R388 of GRP170 active site that contacts lobe Ib of BiP NBD and arginine-arginine stacking arrangement between R66 and R60 from GRP170 and BiP respectively. **c**, displays the interaction between a hook domain helix and BiP lobe IIb with hydrogen bonds highlighted involving N741 and E749 from GRP170. Sequence alignments show the interface residues are well conserved across all GRP170 and Hsp110 chaperones (Hs human, Ms mouse, Dm drosophila, Sc yeast, Pf plasmodium falciparum).

The first interface region is formed through interactions between the NBDs of BiP and GRP170 that are orientated towards each other (Fig. 2a, b). This region features extensive van der Waals and hydrogen bonding contacts. The residues E331, Q329 and N323 at the top of the GRP170 NBD interact with R283, K287 and N82 of BiP. Core interactions are mediated by residues that also form part of GRP 170 ATP active pocket including R388, S321, and A322. These residues engage G58 and Q57 that form part of a loop from lobe Ib of BiP. Additionally, GRP170 residue R66 forms a distinctive arginine–arginine stacking interaction with BiP R60, contributing to the electrostatic stabilization of the complex.

The second interface region involves binding between BiP’s lobe IIb and the hook domain of GRP170 (Fig. 2c). Key residues in this area include GRP170 N741, which forms hydrogen bonds with the side chain of BiP H303 and the main chains of A325 and R324. A salt bridge between GRP170 E749 and BiP K370 further stabilizes the interaction surface. Moreover, GRP170 residues E744, A745, and F748 from one side of a hook domain helix establishes van der Waals contacts with residues A325 and E329 from BiP.

Importantly, nearly all critical interaction residues are conserved across eukaryotic ER-resident GRP170 and cytosolic Hsp110 isoforms. This conservation underscores the evolutionary significance of the BiP–GRP170 interface and highlights its importance across diverse species.

### GRP170 active site and nucleotide asymmetry in the heterodimer complex

The GRP170 active site contains ATP, whereas BiP lacks any bound nucleotide (Fig. 3a, b). An important feature of nucleotide binding in GRP170 ATPase pocket is a hydrogen bond between the N6 atom of the adenine ring and the main chain of Q57 in lobe Ib of BiP NBD. This interaction is facilitated by BiP NDB forming one side of the GRP170 active site, suggesting that ATP binding contributes directly to the hetero dimer interface. This is consistent with previous biochemical data showing that Hsp70-Hsp110 interactions require ATP^19^.

**Fig. 3:**
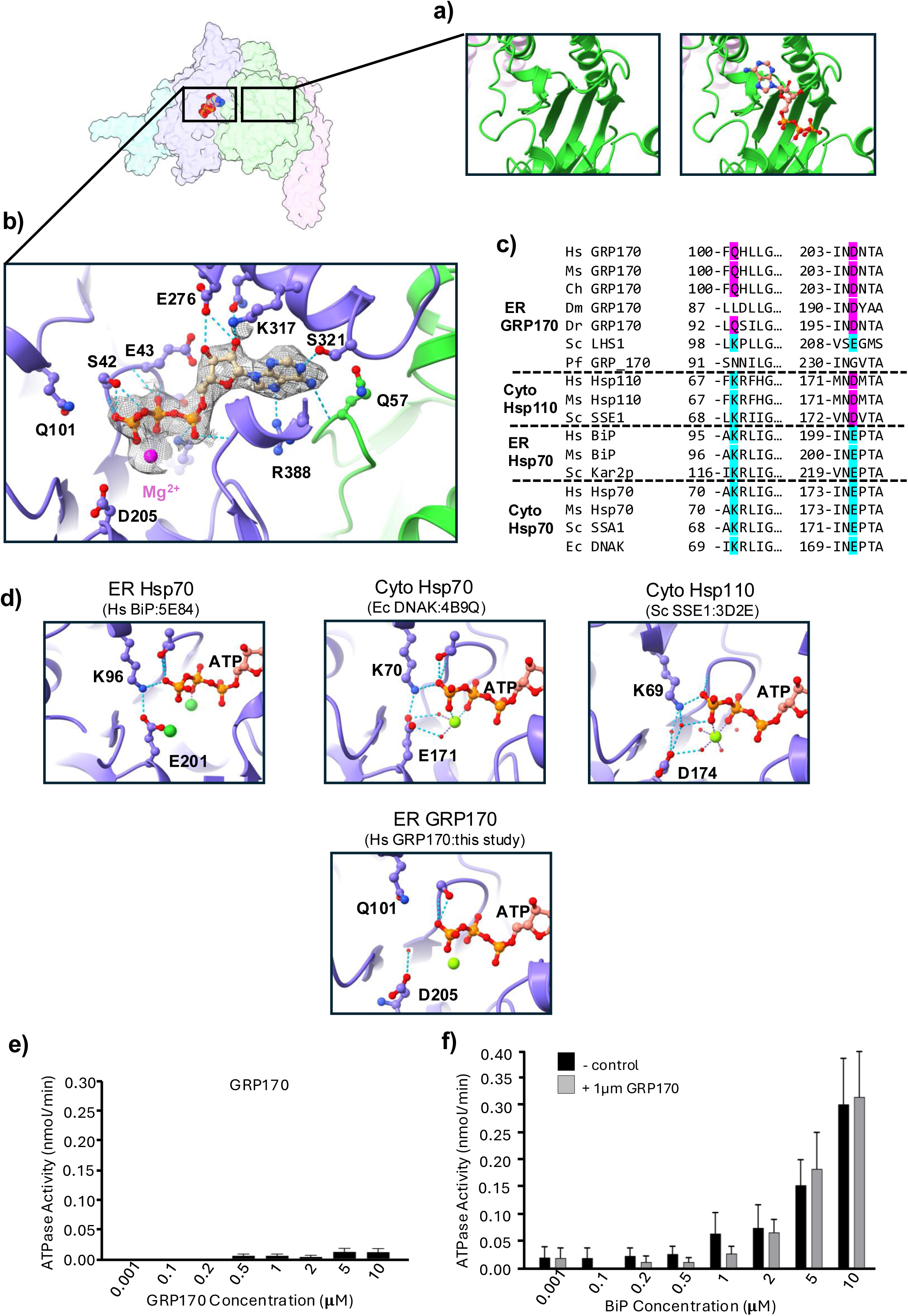
GRP170 active site and the molecular basis for pseudo-ATPase activity. **a**, shows BiP NBD is in apo state in the complex structure, alongside a view of a generated ATP molecule modelled in active site. **b**, ATP molecule in GRP170 active site with corresponding cryo-EM volume in mesh view. Hydrogen bonding between ATP and key residues within the active site are shown, including a Mg^2+^ atom binding to β and γ-phosphates. **c**, Sequence alignment comparing Hsp70 and Hsp110 from both cytosol and ER highlighting the catalytic Lys residue. **d**, a cartoon view of ATP in the active site from cytosolic Hsp70, Hsp110, ER Hsp70 and ER GRP170. The catalytic Lys residue hydrogen bonds to the γ-phosphate orientating it for catalysis. However, in GRP170 the Lys is substituted to Gln which is too far to hydrogen bond to the γ-phosphate. Not all the hydrogen bonds from the γ-phosphate three oxygens are shown to aid clarity. e, NADH coupled assay measuring ATPase activity of GRP170 sample and when added to recombinantly expressed BiP (error bars are mean ± sem, n=3, two-way Anova test, p= 0.7525 = ns, no significance).

Additional interactions within the ATP-binding pocket include R388 and S321, which engage the adenine ring. Residues E276, N314, and K317 form hydrogen bonds with the ribose moiety. Meanwhile, S42, E43, S44, K46 and G236 mediate both main-chain and side-chain interactions with the phosphate groups. Coordination with Mg^2+^ at the β-and γ-phosphates further stabilizes nucleotide binding (Fig. 3b).

### GRP170 acts as a pseudo-ATPase chaperone

All Hsp70 chaperones possess an essential catalytic lysine residue within the ATPase pocket^20^. This lysine interacts with the γ-phosphate of ATP, positioning it optimally for hydrolysis. Additionally, a salt bridge interaction is formed between the lysine and a glutamic acid that further stabilises the lysine— γ-phosphate hydrolytic configuration (Fig. 3c, d). In cytosolic Hsp110, the glutamic acid is substituted to an aspartate but still hydrogen bonds to the catalytic lysine residue via water molecules to maintain the critical lysine— γ-phosphate interaction.

In nearly all GRP170 homologs the catalytic lysine is substituted for a glutamine. Inspection of our structure shows that the shorter side chain of Q101 residue is unable to reach and interact with the γ-phosphate or D205, disrupting the water mediated hydrogen bonding network (Fig. 3d). Furthermore, visualising ATP (and not ADP) in the active site also suggests an inactive ATPase. Therefore, our structure suggests that lysine to glutamine substitution would dramatically reduce hydrolysis.

To test this notion, we measured the ATPase activity of our GRP170 complex sample alone and in a mixture with recombinantly purified BiP. We observed very low activity in GRP170 sample, and no effect on recombinant BiP activity when GRP170 sample was added (Fig. 3e). Thus, GRP170 acts as a pseudo-ATPase chaperone, retaining nucleotide-binding capacity but lacking efficient catalytic activity.

### Hook domain motion is coupled to NBD and β-sandwich domain

To assess structural flexibility in our cryo-EM data, we performed 3D Flexible refinement (3DFlex) analysis^21^, which revealed three regions of motion within the canonical density: the hook, BiP NBD and GRP170 β-sandwich domain (Extended Data Fig. 5, Extended Data Movie 1-4). The hook exhibited the greatest flexibility, shifting 19 Å towards the β-sandwich domain and the putative BiP SBD (Extended Data Movie 2), and showed 26 Å lateral movement relative to BiP and GRP170 NBD Extended Data Movie 4). Notably, motion in the hook appeared to be coupled to subtle movements within the BiP NBD that tended towards a more compact BiP NBD, which was further linked to motion in the β-sandwich domain. These motions are based on apo BiP interacting with GRP170-ATP from our cryo-EM data. A comparison of static structures with BiP bound to different nucleotides indicates a change in the position of lobe IIb that further restricts NBD, which in turn facilitates ATP binding (Extended Data Fig. 6), and shows that apo BiP NBD closely resembles the Hsp70 NBD-ADP state.

## Discussion

Our structural analysis of GRP170-ATP-BiP isolated from HEK293 cells reveals novel insights into the architecture of a critical ER chaperone complex and provides a rationale to previous biochemical/cellular data, that collectively suggest a new paradigm of how GRP170-BiP may engage misfolded protein substrate.

The salient feature of the structure is the α-helical domain adopting a striking hook shape that folds back towards the GRP170 NDB. Previous cellular studies have shown that the α-helical domain of GRP170 is able to engage misfolded substrate^3^. However, it was unclear how this was linked to the canonical SBD/β-sandwich domain as crystal structures of yeast Hsp110 indicated that the domains were far apart and orientated away from each other. Our structure shows that the hook domain bends towards the BiP SBD, possibly even making contact. Furthermore, flexibility analysis indicates that the hook domain undergoes significant motion that is linked to more subtle movements in both BiP NDB and the β-sandwich domain. These observations, together with earlier cellular analysis, suggest that GRP170 hook domain may collaborate with the BiP SBD to clamp onto substrate protein (Fig. 4).

**Fig. 4:**
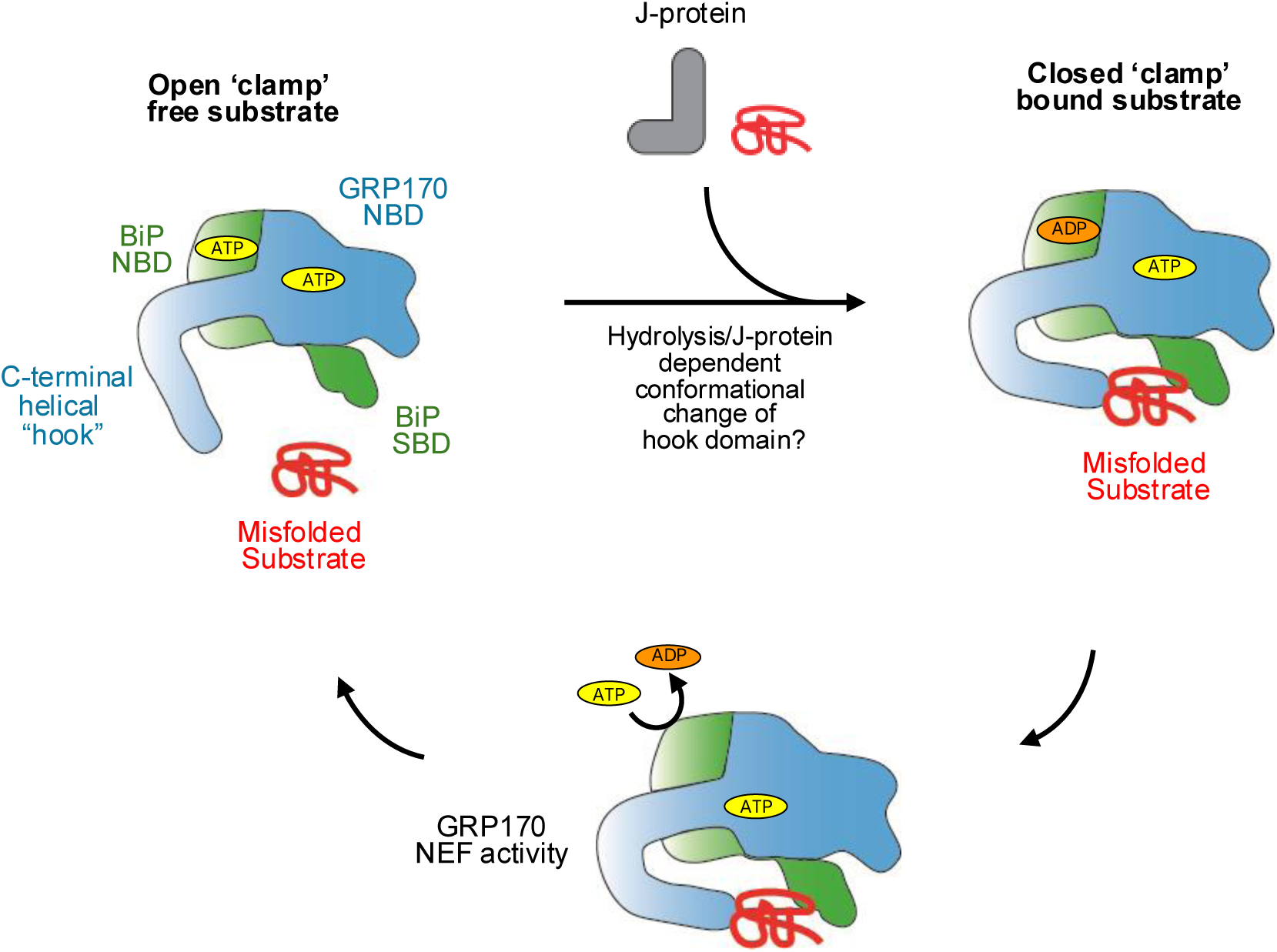
A clamp model for GRP170-BiP association with substrate. GRP170 forms a stable heterodimer with BiP in the presence of ATP. Substrate recruitment is mediated by a J-domain protein, which triggers ATP hydrolysis in BiP. This induces conformational changes in the hook domain, enabling it to clamp the substrate in coordination with BiP SBD. Subsequent ADP exchange, facilitated by lateral hook domain movement, promotes substrate release and resets the complex to its open state.

Interestingly, GRP170 acts as a pseudo-ATPase chaperone: it can bind ATP but is unable to hydrolyse it. To our knowledge, this is the first example of pseudo-ATPase regulation in an ATP-binding chaperone. This finding explains how GRP170 can associate with substrates in an ATP independent manner^3^. This property may serve two important roles. First, it aids stabilisation of the GRP170 NBD by contributing towards formation of the complex and providing a level of regulation by favouring heterodimer association in the presence of ATP. Second, it enables GRP170 to act as a scaffold whilst operating as a NEF in response to BiP ATP hydrolysis cycle, which would also support BiP SBD dependent movements. This may only be feasible if GRP170 itself does not undergo allosteric changes induced by ATP hydrolysis.

Current models suggest that GRP170/Hsp110 transiently interacts with BiP/Hsp70 serving as a NEF in a defined step that facilitates ADP to ATP exchange thereby promoting the transition of Hsp70 from closed to open state. However, the structure of the heterodimer complex shows a substantial interaction surface that is supported by numerous van der Waals and hydrogen bond contacts. Moreover, it purifies as an ER chaperone complex from cells unlike cytosolic Hsp90 chaperones, suggesting against a transient interaction.

Once ATP is bound to GRP170 it contributes to the heterodimer complex formation resulting in a buried and inaccessible active site that is unable to hydrolyse ATP. In contrast, BiP active site is readily accessible for nucleotide exchange facilitated by GRP170 binding at the interface 2 which causes displacement of the lobe IIb, with subtle movements in lobe Ib.

ATP hydrolysis within BiP NBD active site could be coupled to GRP170 ability to interact with misfolded substrate via lobe IIb displacement causing movement of the hook domain relative to putative position of BiP SBD. The opening and closing of the clamp, which would enable substrate engagement, could be linked to nucleotide hydrolysis and exchange activity within BiP NBD. In this model, both GRP170 and BiP form a stable heterodimer complex that combines GRP170 NEF activity and chaperone function in collaboration with BiP as a single complex molecular chaperone unit.

## Methods

### Expression and purification of GRP170-ATP-BiP complex from HEK cells

A full-length construct of human GRP170 containing a triple FLAG tag positioned between the signal sequence (1-32aa) and the mature protein (33-999), was inserted into pcDNA3.1+ vector for expression in Expi293F cells. 24h before transfection the cells were diluted to a density of 1.5 × 10^6^ per ml. Transient transfection was performed at a cell density of 2.5 × 10^6^ per ml using polyethylenimine (PEI). 1mg/ml of plasmid DNA was added to PEI at a ratio of 1:3 in Opti-MEM® medium (Thermo-Fisher Scientific) and incubated at room temperature for 10 min, then the mixture was added to cells. 24h post-transfection, valproic acid was added to the cells at a final concentration of 3.2 mM. The cells were harvested 48h post-transfection by centrifugation at 700g for 10 min at 4°C. Pellets were resuspended in buffer A (50 mM Hepes pH 7.5, 150 mM NaCl, 5 mM MgCl_2_, 5 mM CaCl_2_, 1% Triton-X) that was supplemented with DNase (Sigma), Benzonase (Merck) and EDTA-free protease inhibitor tablet (Merck) and lysed by passing through Dounce homogeniser. Lysate was clarified by centrifugation for 1 hour at 18500 x g at 4°C. The supernatant was incubated with anti-Flag®M2 Affinity resin (Sigma-Aldrich) overnight at 4°C and then transferred to gravity flow column and washed with buffer B (50 mM Hepes pH 7.5, 150 mM NaCl, 5 mM MgCl_2_, 5 mM CaCl_2_), followed by a high salt wash buffer C (buffer B but with 1 M NaCl) and then incubated for 15 minutes and washed with ATP buffer D (buffer B with 1 mM ATP), before eluting with buffer E (buffer B with Flag peptide). Eluted sample was then pooled, concentrated and applied to size-exclusion chromatography performed using a superdex 200 10/300 GL column in buffer F (buffer B with 1 mM DTT). GRP170 bound to endogenous BiP sample resolved clearly away from the void peak (Extended Data Fig. 1), was pooled and concentrated for further analysis.

### Grid preparation

UltraAuFoil R 2/2, 300-mesh grids were glow-discharged in a Fischione Model 1070 NanoClean at 70% power for 2 min 30 s. Purified sample at 2.5 mg ml⁻ ¹ in SEC buffer F was applied (3 µl) to each grid in a Vitrobot Mark IV (Thermo Fisher Scientific; 4 °C, 95% RH), blotted for 7 s with blot force −2 and plunge-frozen into liquid ethane. Grids were screened on a Glacios (Thermo Fisher) for ice thickness and sent to a Titan Krios for data collection.

### Cryo-EM data acquisition

Data were collected on Titan Krios G3 microscope operated at 300 kV with field-emission gun. The microscope was equipped with a BioQuantum energy filter (slit width 20 eV) and a Gatan K3 detector in super-resolution mode. Automated acquisition used EPU in Faster Acquisition mode. 15266 movies were recorded and binned 2X in EPU to yield a final pixel size of 0.825 Å at a magnification of 105,000X. Movies contained 60 fractions at ∼1.0 e⁻ Å² per fraction for a total exposure of 59.98 e⁻ Å². The defocus range was set between −0.7 and −2.2 µm. Data collection statistics are listed in Extended data Table 1.

### Image processing

Movies were motion corrected using patch motion correction and CTF parameters were estimated with Patch CTF in cryoSPARC (v4.7)^22^. Further downstream processing was also conducted in cryoSPARC (v4.7). An initial 2,786,909 particles were blob-picked to generate templates for Topaz training^23^. Automated particle picking with the trained Topaz model was followed by one round of reference-free 2D classification, yielding 482,963 particles for ab initio reconstruction and non-uniform (NU) refinement. These particles were then used to retrain Topaz and additional rounds of 2D and 3D classification were performed. A third Topaz training recentred on flexible domain, with merging of previous datasets and removal of duplicates, resulting in 598,978 particles. Heterogeneous refinement separated conformational classes, and the best class was refined with NU-refinement, reference-based motion correction, global and local CTF refinement. Additional 3D classification and focused refinements with soft masks further improved domain resolution and density. Final reconstructions reached 2.7 Å resolution (gold-standard FSC 0.143 criterion). Post-processing and map sharpening were performed in cryoSPARC with DeepEMhancer^24^. The cryo-EM processing workflow is shown in Extended Data Fig. 2.

The structure was further analysed with 3DVA^25^ and 3DFlex^21^ to assess conformational heterogeneity in cryoSPARC v4.7. For 3DVA, a mask was created that focused on the BiP SBD. The data filter was set at a low resolution of 6-8 Å and a cluster was reconstructed using non-uniform refinement. For 3DFlex, the particles were prepared for training in flex data prep. The mesh was generated with 40 tetrahedra per cell, covering the entire structure, and a rigidity weight of 0.15. The training step was performed with latent centring strength of 0.2, rigidity (λ) of 0.007, and three latent dimensions.

### Model building and refinement

AlphaFold 3^26^ prediction was used to generate an initial model for GRP170 that was rigid body fitted into cryo-EM density using UCSF ChimeraX v1.1^27^ and then rigid body refined in Phenix v1.21^28^. For BiP density, Hsp70 NBD (3D2E) was used as an initial model, with lobe IIb requiring significant rebuilding. For the overall complex structure, iterative model rebuilding and real space refinement using secondary structure, rotamers and Ramachandran restraints was performed in Coot v0.9.8.6^29^ and Phenix. Model geometry was validated with MolProbity in Phenix. Figures were prepared using UCSF ChimeraX v1.1.

### Recombinant BiP expression and purification

A human BiP construct (28-654) containing a N-terminal His and Strep tag was inserted into pET 17b vector for expression in E. coli BL21 (DE3) cells as previously described in Kopp et al^30^. Briefly, cells were grown in auto-inducing media (Formedia) at 37°C for 3 hours and then overnight at 22°C. His-tag purification was conducted using HisPur cobalt resin (ThermoFisher Scientific), with several wash steps starting with buffer G (50 mM Hepes pH 7.3, 400 mM NaCl, 10 mM MgCl_2_, 10 mM Imidazole) followed by high salt wash (buffer G but with 1M NaCl), and then final wash buffer H (50 mM Hepes pH 7.3, 150 mM NaCl, 10 mM MgCl_2_, 20 mM Imidazole). Protein was eluted with high imidazole buffer I (50 mM Hepes pH 7.3, 150 mM NaCl, 10 mM MgCl_2_, 350 mM Imidazole). Eluted protein was pooled and then incubated with Step-TactinXT resin (IBA life sciences) that was washed with buffer J (50 mM Hepes pH 7.3, 150 mM NaCl, 10 mM MgCl_2_, 1 mM EDTA) and eluted with buffer K (50 mM Hepes pH 7.3, 150 mM NaCl, 10 mM MgCl_2_, 1 mM EDTA, 50 mM biotin), pooled and concentrated for further analysis.

### NADH-couple ATPase assay

Reactions were conducted in 50 mM Hepes pH 7.5, 40 mM KCl, 4 mM MgCl_2_, 2 mM PEP, 1 mM ATP, 1% pyruvate kinase/lactic dehydrogenase buffer, using an optical bottom 96 well plate (thermo scientific) at 37°C. NADH standards were measured at a range of 0-400 µM concentration. The reaction samples were 100µl in volume and contained 300 µM NADH. Absorbance of 340 nM was recorded in 90-second cycles using a CLARIOstar plate reader (BMG Lab tech). NADH standards curves enabled calculation of NADH depletion rates, which were converted to ATP hydrolysis rate after correcting for controls.

### Peptide mass fingerprinting

Samples were run on SDS-PAGE gel and stained with Coomassie, with specific bands excised out. Bands were subjected to in gel tryptic digestion and MALDI mass spectrometry at the Proteomics facility (University of St Andrews, UK) for protein ID services.

## Extended Data Figure legends

**Extended Data Fig. 1.**
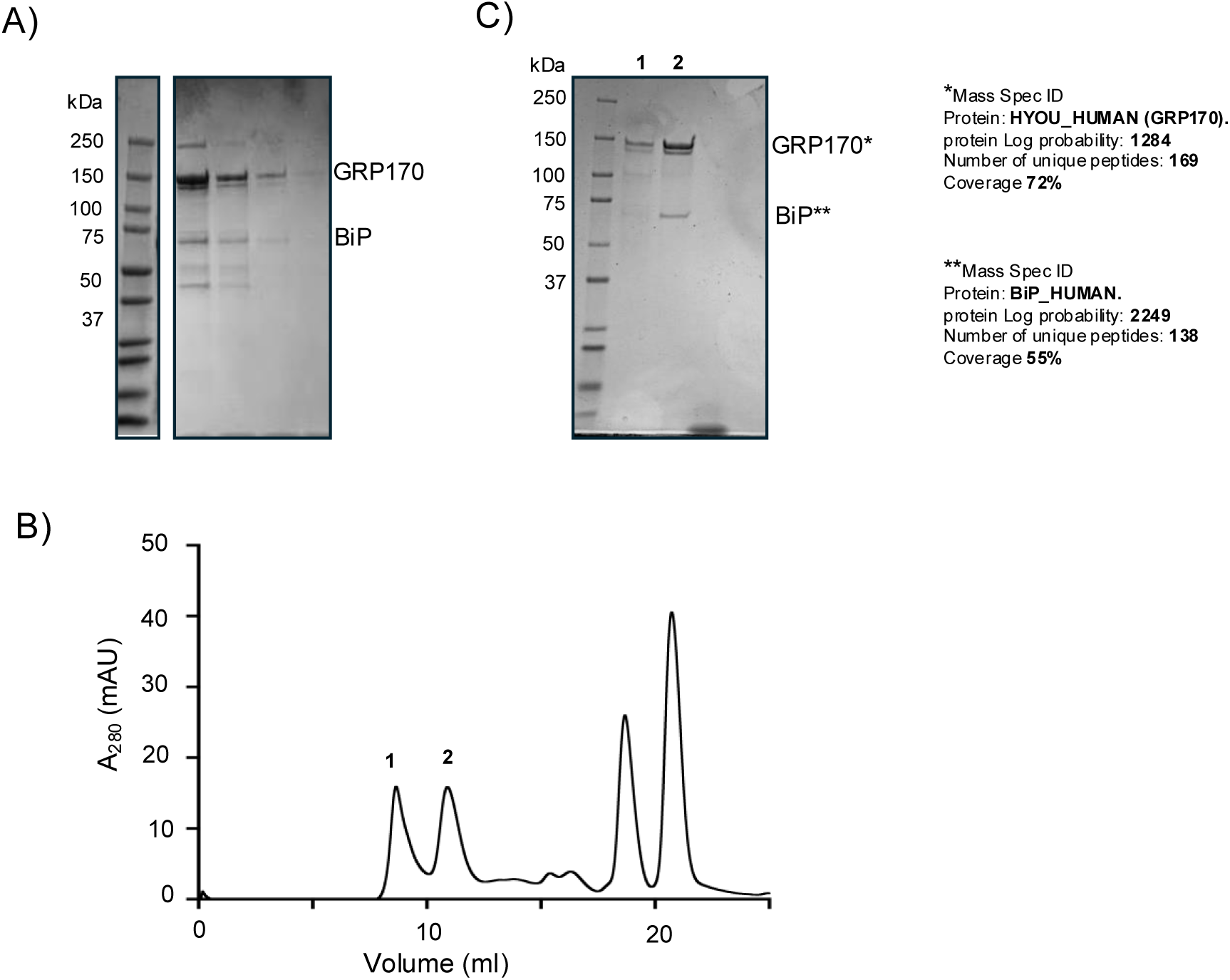
Purification of chaperone complex. a) An SDS-PAGE gel showing the eluted fraction after FLAG-GRP170 affinity purification step from Expi293 cell lysate, stained with Coomassie blue. b) A size exclusion chromatogram of GRP170-BiP complex after FLAG affinity step. c) SDS-PAGE gel showing peaks 1 and 2 from size exclusion chromatograph along with band identification by mass spectrometry.

**Extended Data Fig. 2.**
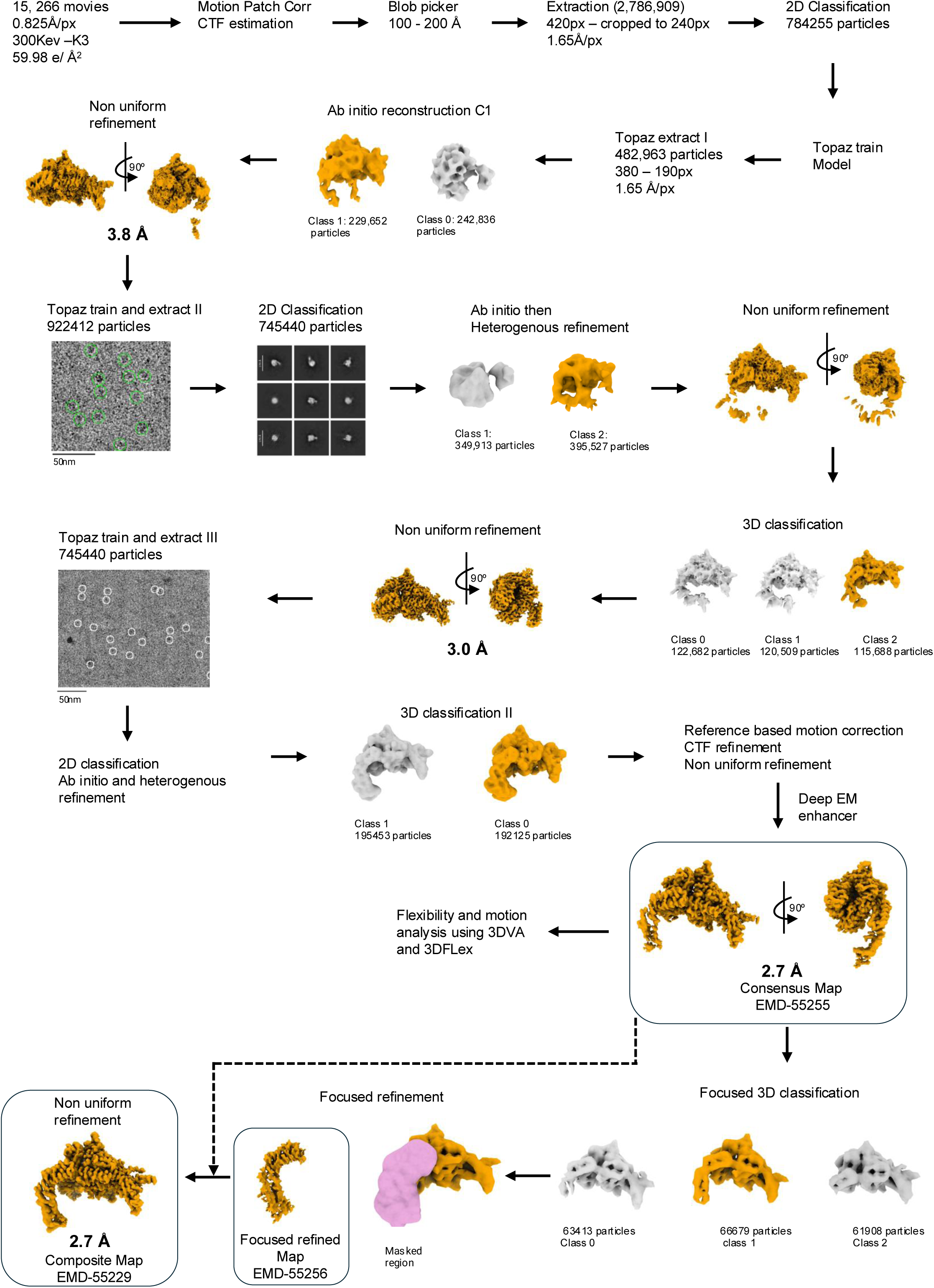
Cryo-EM data processing workflow. Data processing pipeline highlighting each stage towards generating high resolution maps.

**Extended Data Fig. 3.**
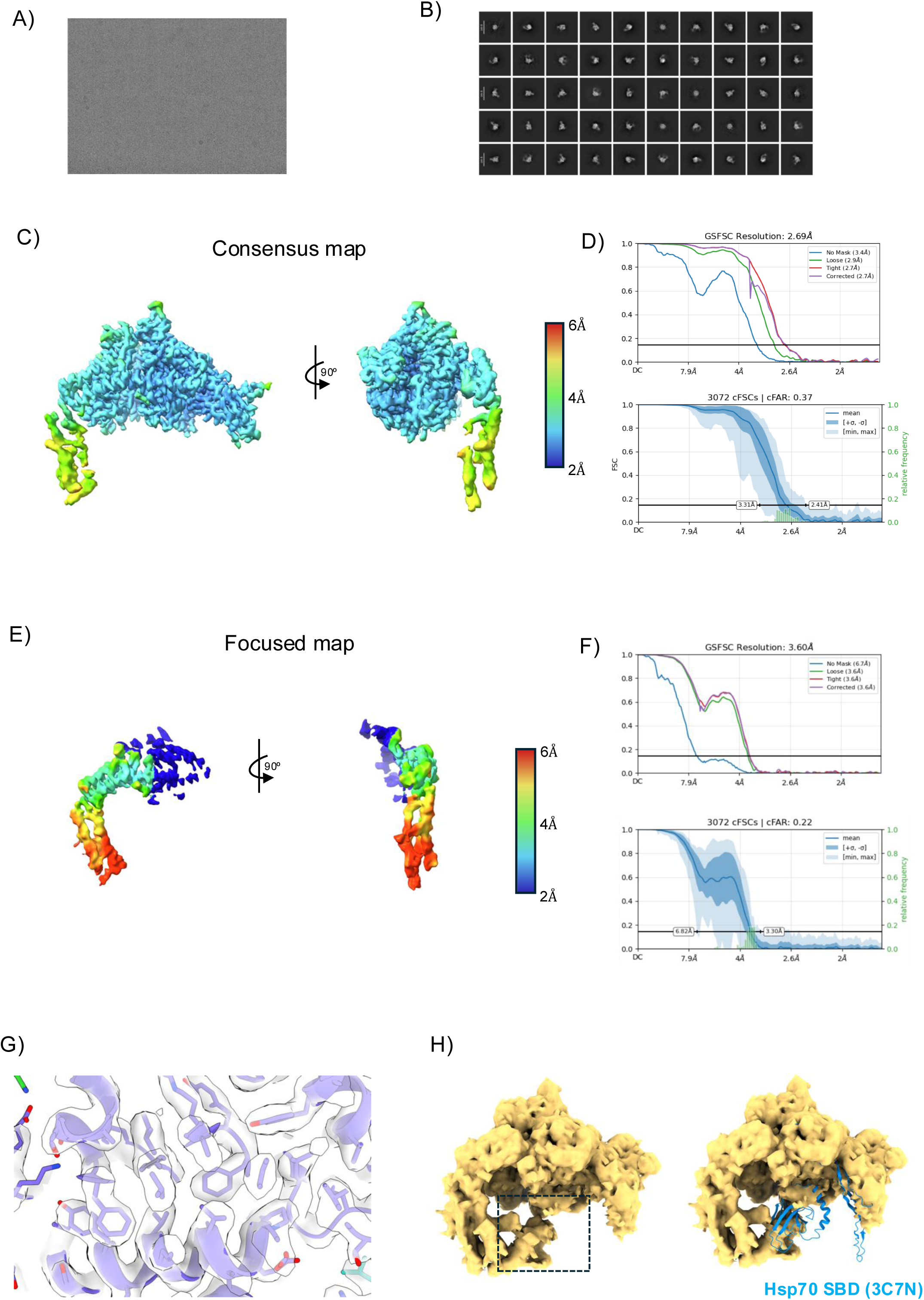
Local resolution and orientation analysis. a) A micrograph containing protein sample. b) 2D class averages showing different orientations of our protein complex. c) Local resolution estimate of consensus map. d) GSFSC curve and conical FSC area ratio (cFAR) indicating the global resolution and a measure of orientation bias in consensus map. e) Local resolution estimate of focused map. f) same as (d) but for focused map. g) illustrates a cross section of the density map corresponding to the GRP NBD with fitted atomic model. h) 3DVA identifies a structural cluster that potentially shows the BiP SBD in contact with hook domain when contoured at low threshold and low resolution (highlighted by dashed box). SBD from 3C7N modelled into density suggests that the SBD adopts a significantly different position or conformation in the cryo-EM map.

**Extended Data Fig. 4.**
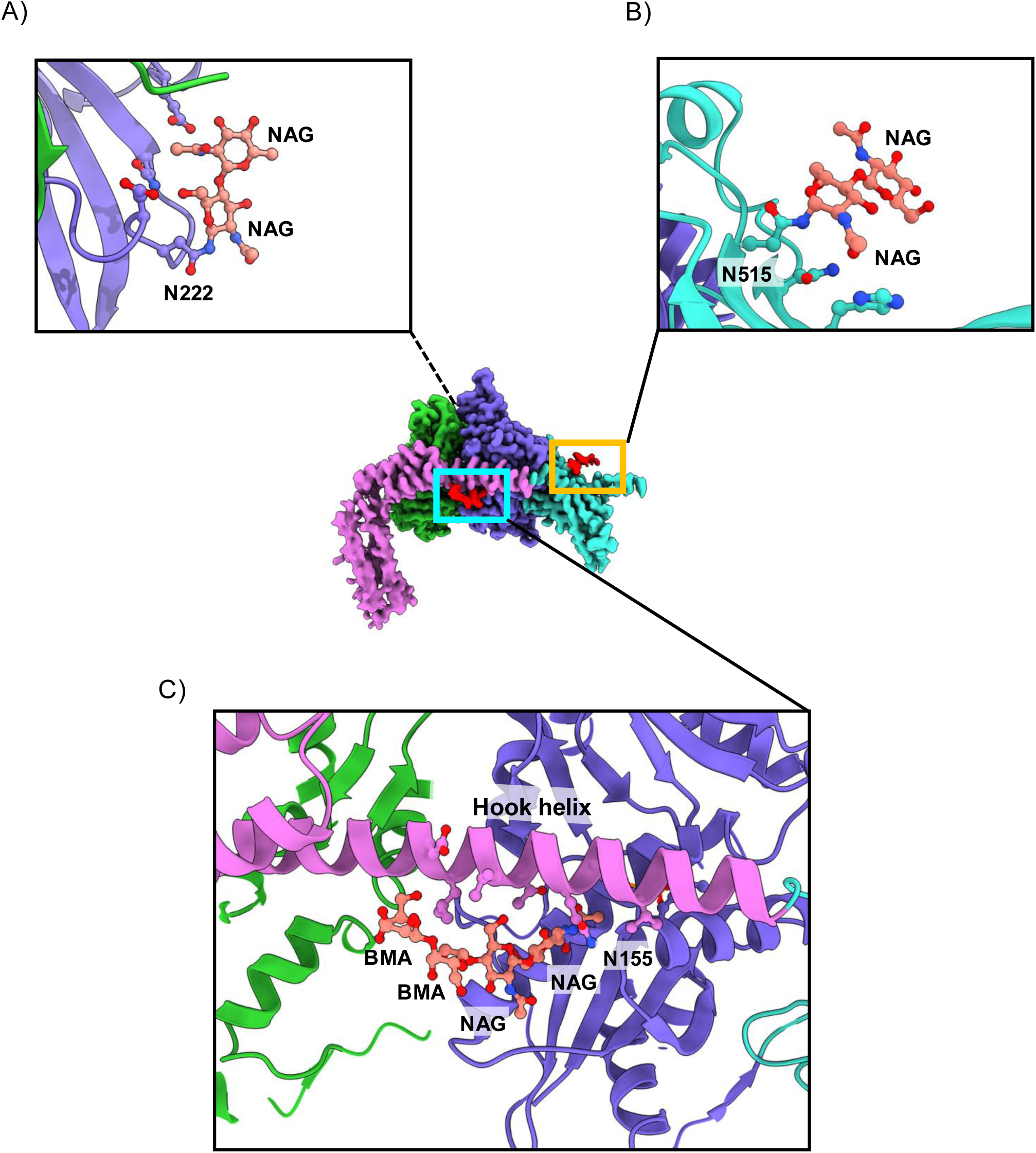
Glycosylation site supports hook domain extension. Three N-linked glycans were modelled onto GRP170: a) N222, b) N515, c) N155. The glycan at N155, located within GRP170 NBD, appears to stabilise hook domain extension through van der Waals interactions.

**Extended Data Fig. 5.**
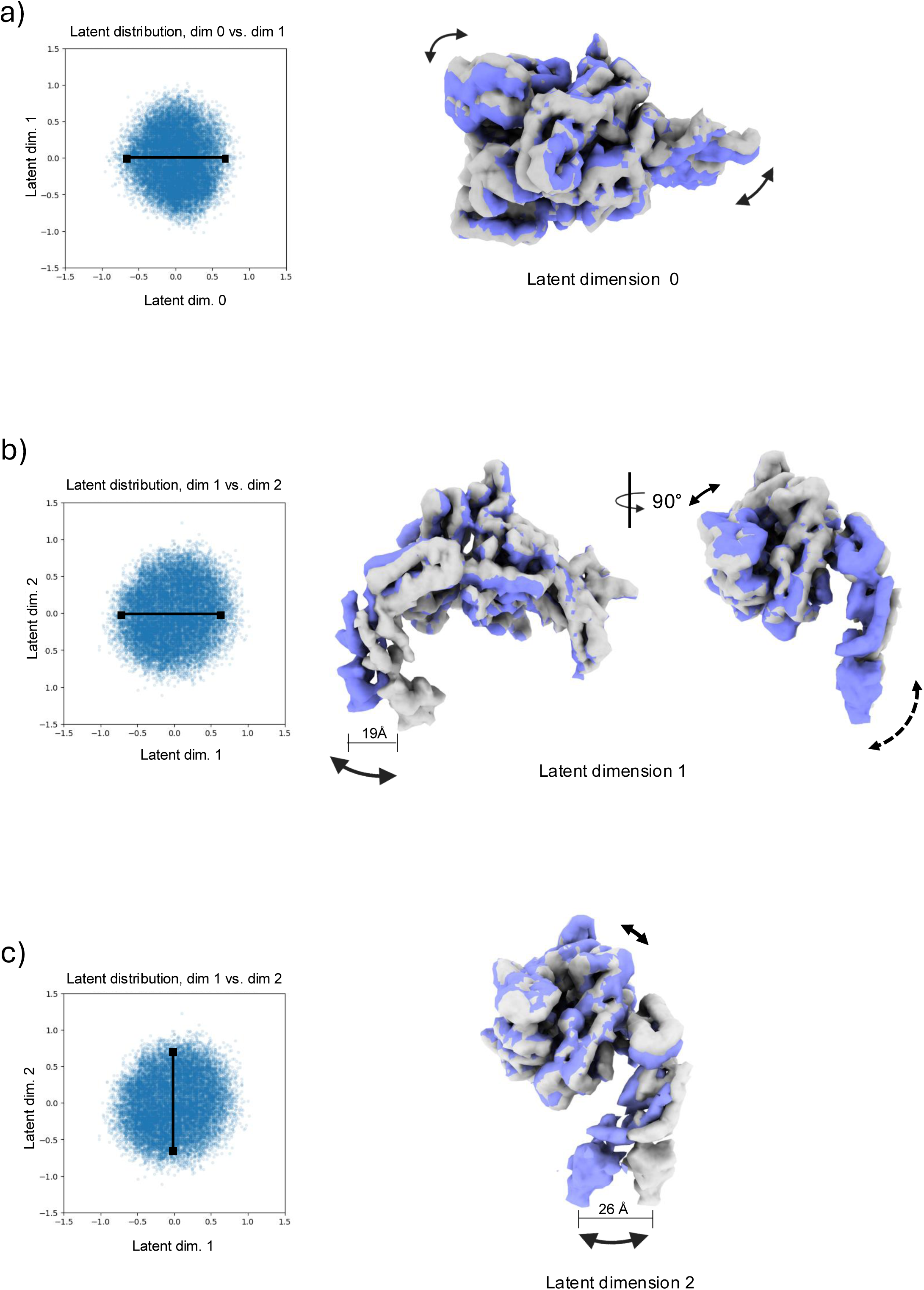
Flexibility and motion analysis. Convected densities generated from 3DFlex at -0.7 (grey) to +0.7 (purple) standard deviations in latent space along a specific coordinate axis (dimension) as shown in the corresponding scatter plot. a) Dimension 0 -A top view shows motion within the GRP170 b-sandwich and BiP NBD lobe Ib. b) dimension 1-A front and side view captures the motion of the hook domain that seems to be coupled to subtle movement in lobe Ib that restricts BiP NBD. The dashed line indicates movements perpendicular to plane. c). dimension 2 captures the lateral motion of the hook domain.

**Extended Data Fig. 6.**
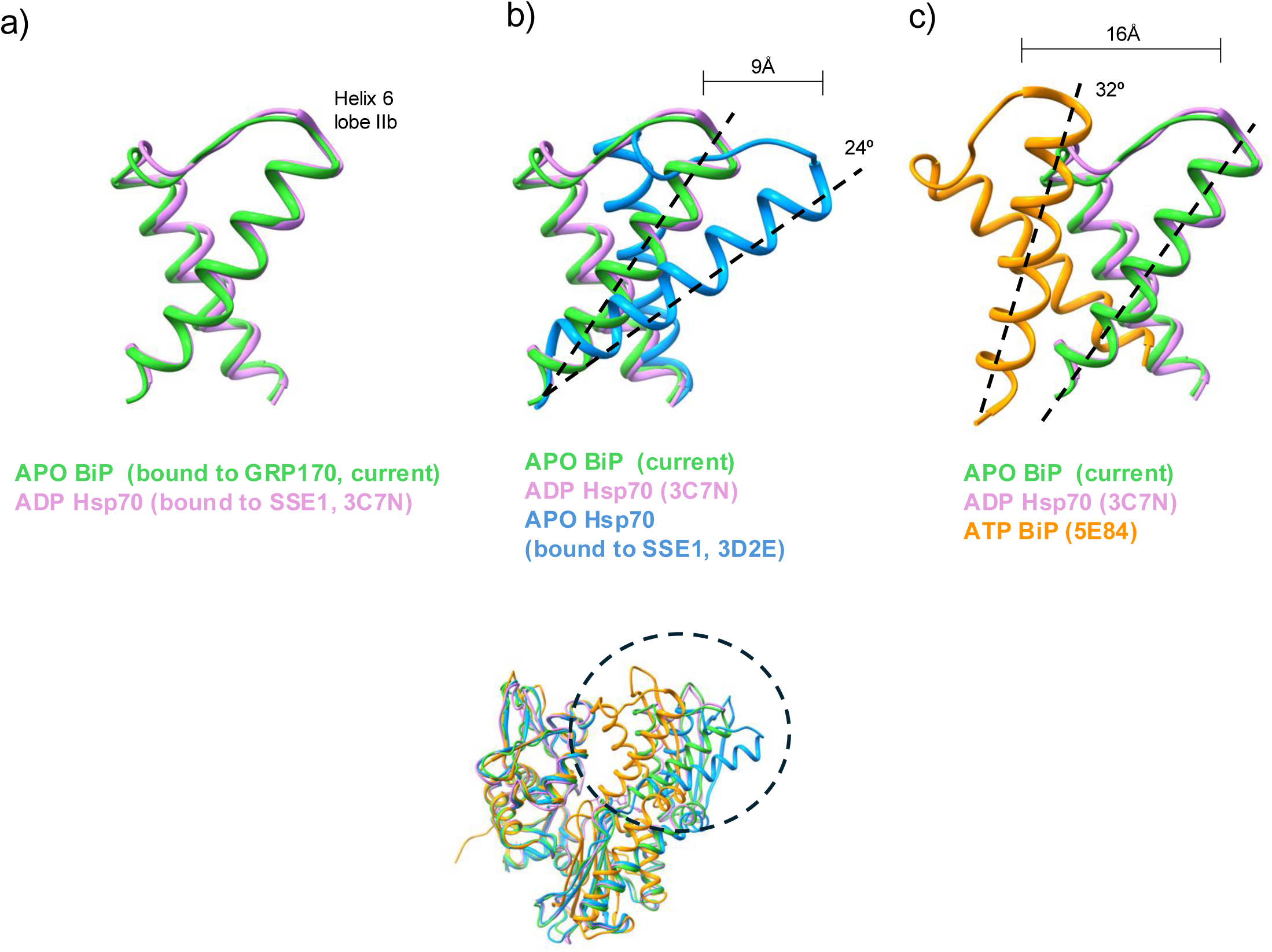
A comparison of lobe IIb in various nucleotide bound states. a) Helix 6 and 7 of lobe IIb from apo BiP bound to GRP170 closely resembles the ADP Hsp70 NBD bound to Sse1. b) Helix 6 from apo Hsp70 bound to Sse1 shows significant deviation to both the apo BiP and ADP bound Hsp70 structures. c) same as b) but with ATP bound Hsp70 NBD.

**Extended Data Table 1.**
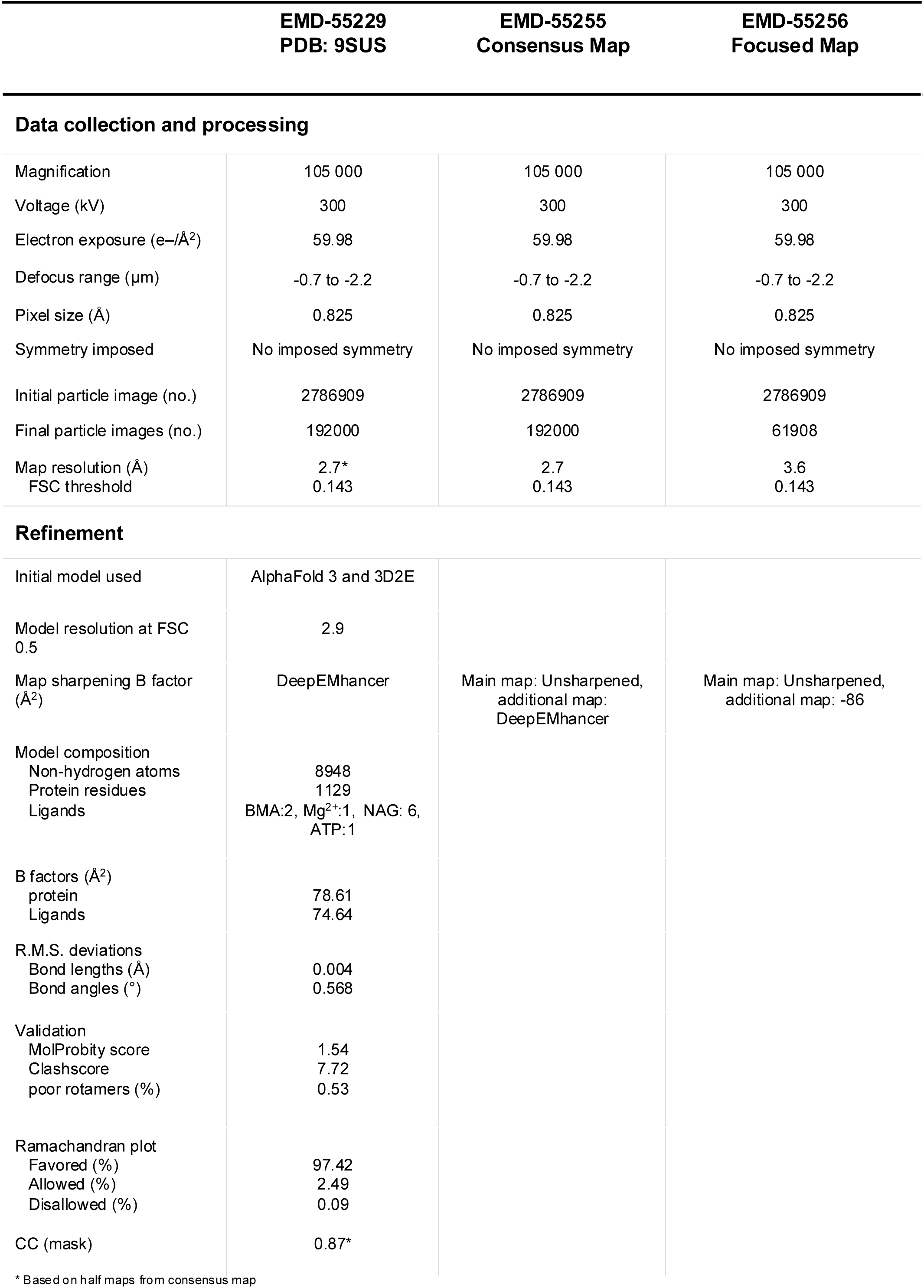
Cryo-EM data collection, refinement and validation statistics.

## Author contributions

YBK: Conceptualisation, data curation, formal analysis, investigation, methodology, validation, visualisation, writing-review and editing. KW: Formal analysis, investigation, reviewing final draft. SB: reviewing final draft. MMUA: Conceptualisation, data curation, formal analysis, funding acquisition, project administration, supervision, validation, visualization, writing-original draft, writing-review and editing.

## Data availability

All data will be available upon publication. Cryo-EM maps have been deposited in the Electron Microscopy Data Bank (EMDB) with the following accession numbers: EMD-55229 for GRP170-ATP-BiP composite map, EMD-55255 for unsharpened consensus map, EMD-55256 for focus refined map of hook domain. The atomic model has been deposited in the protein data bank (PDB) with accession code: 9SUS. Source data are provided with this paper.

## Acknowledgements

We thank: former and current members of the Ali lab including Piotr Nowak and Archna Shah; Harry Low and Tiago Costa for useful discussions; the Mass spectrometry facility at St Andrews for protein ID services; the Centre of Structural Biology, including Paul Simpson; Suhail Islam for IT support; the cryo-EM computing platform at Imperial College funded by a BBSRC equipment grant (BB/X019284/1); Diamond for access and support for cryo-EM facilities at eBIC, proposal BI39228; the Department of Life Sciences for funding of a PhD studentship (Widening Participation scheme).

