## Supplementary information for "Structure of ER chaperone complex GRP170-ATP-BiP suggests a new model for substrate engagement"

Supplementary Information - Sequence alignment of eukaryotic GRP170 and Hsp110

```
sp|Q9Y4L1|/1-999     1 MADKVRRQRPRRRVCWALVAVLLADLLALSDTLAVMSVDLGSESMKVAIVKPGVPMEIVLNKESRRKTPVIVTLKE---------  76
sp|Q9JKR6|/1-999     1 MAATVRRQRPRRLLCWALVAVLLADLLALSDTLAVMSVDLGSESMKVAIVKPGVPMEIVLNKESRRKTPVTVTLKE---------  76
sp|Q60432|/1-999     1 MAATVRRQRPRRLLCWTLVAVLLADLLALSDTLAVMSVDLGSESMKVAIVKPGVPMEIVLNKESRRKTPVTVTLKE---------  76
tr|O46067|/1-923     1 MKLVL------------LLSALLA-GIALSQGAAVMSVDLGSEWMKVGVVSPGVPMEIALNRESKRKTPAILAFRD---------  63
sp|P36016|/1-881     1 MRNVLR-------------LLFLTAFVAIGSLAAVLGVDYGQQNIKAIVVSPQAPLELVLTPEAKRKEISGLSIKRLPGYGKDDP  72
sp|Q7ZUW2|/1-980     1 MREKL--------SLWAIFCLVVAFLPSQTESVAVMSVDLGSEWMKVAIVKPGVPMEIVLNKESRRKTPVAVCLKE---------  68
tr|C0H5H0|/1-932     1 MRPRF--------FLFLLFIIYIYNSLRI-KCSSLLGIDFGNEYIKVSIVSPGKGFNILLNNQSKRKITNSISFAN---------  67
sp|Q92598|/1-858     1 M--------------------------------SVVGLDVGSQSCYIAVARAG-GIETIANEFSDRCTPSVISFGS---------  43
sp|Q61699|/1-858     1 M--------------------------------SVVGLDVGSQSCYIAVARAG-GIETIANEFSDRCTPSVISFGS---------  43
sp|P32589|/1-693     1 MS-------------------------------TPFGLDLGNNNSVLAVARNR-GIDIVVNEVSNRSTPSVVGFGP---------  44

sp|Q9Y4L1|/1-999    77 --NERFFGDSAASMAIKNPKATLRYFQHLLGKQADN-------PHVAL--------------------YQARFPEHELTFDPQRQ 132
sp|Q9JKR6|/1-999    77 --NERFLGDSAAGMAIKNPKATLRYFQHLLGKQADN-------PHVAL--------------------YRSRFPEHELIVDPQRQ 132
sp|Q60432|/1-999    77 --NERFLGDSAAGMAIKNPKATLRYFQHLLGKQADN-------PHVAL--------------------YRDRFPEHELNIDPQRQ 132
tr|O46067|/1-923    64 --GTRTIGEDAQTIGIKDPNSAYGYLLDLLGKTIDN-------PIVDL--------------------YRKRFPYYNIVGDPERN 119
sp|P36016|/1-881    73 NGIERIYGSAVGSLATRFPQNTLLHLKPLLGKSLEDE------TTVTL--------------------YSKQHPGLEMV-STNRS 130
sp|Q7ZUW2|/1-980    69 --NERLFGDGALGVAVKNPKVVYRFLQSILGKTADN-------PQVAE--------------------YQKHFPEHQLQKDEKRG 124
tr|C0H5H0|/1-932    68 --KFRTYDEESKIYSTKYPQLTLLNSNNILGYNLFDSLKNKENFVIENYDENNEEFYSDINNYDFSNDFGSKYYSYDYVVDHKRG 150
sp|Q92598|/1-858    44 --KNRTIGVAAKNQQITHANNTVSNFKRFHGRAFND-------PFIQK--------------------EKENLSYDLVPLKNGGV  99
sp|Q61699|/1-858    44 --KNRTIGVAAKNQQITHANNTVSSFKRFHGRAFND-------PFIQK--------------------EKENLSYDLVPMKNGGV  99
sp|P32589|/1-693    45 --KNRYLGETGKNKQTSNIKNTVANLKRIIGLDYHH-------PDFEQ--------------------ESKHFTSKLVELDDKKT 100

sp|Q9Y4L1|/1-999   133 TVHF-QISSQLQFSPEEVLGMVLNYSRSLAEDF-------AEQPI---------KDAVITVPVFFNQAERRAVLQAARM-AGLKV 199
sp|Q9JKR6|/1-999   133 TVRF-QISPQLQFSPEEVLGMVLNYSRSLAEDF-------AEQPI---------KDAVITVPAFFNQAERRAVLQAARM-AGLKV 199
sp|Q60432|/1-999   133 TVRF-QISPQLQFSPEEVLGMVLNYSRSLAEDF-------AEQPI---------KDAVITVPAFFNQAERRAVLQAARM-AGLKV 199
tr|O46067|/1-923   120 TVVF-RKSDTDEFSVEELVAQLLVKAKQFAQES-------VQQPI---------TECVLTVPGYFGQAERREALLSAAQL-ANLKV 186
sp|P36016|/1-881   131 TIAF-L-VDNVEYPLEELVAMNVQEIANRANSLLKDRDARTEDFV---------NKMSFTIPDFFDQHQRKALLDASSITTGIEE 204
sp|Q7ZUW2|/1-980   125 TVYF-KFSEEMQYTPEELLGMILNYSRTLAQDF-------AEQPI---------KDAVITVPAYFNQAERRAVLQAAHI-AGLKV 191
tr|C0H5H0|/1-932   151 TINI-KLKDNMVISSEEVTANILGYIKKLAYTH-------LNIDYKVKRNINLNIGCVISVPCNFSQRKKQALINASKI-AGLEL 226
sp|Q92598|/1-858   100 GIKVMYMGEEHLFSVEQITAMLLTKLKETAENS-------LKKPV---------TDCVISVPSFFTDAERRSVLDAAQI-VGLNC 167
sp|Q61699|/1-858   100 GIKVMYMDEEHFFSVEQITAMLLTKLKETAENN-------LKKPV---------TDCVISVPSFFTDAERRSVLDAAQI-VGLNC 167
sp|P32589|/1-693   101 GAEVRFAGEKHVFSATQLAAMFIDKVKDTVKQD-------TKANI---------TDVCIAVPPWYTEEQRYNIADAARI-AGLNP 168

sp|Q9Y4L1|/1-999   200 LQLINDNTATALSYGVFRRKDIN--TTAQNIMFYDMGSGSTVCTIVTYQMVKTKEA-GMQPQLQIRGVGFDRTLGGLEMELRLRE 281
sp|Q9JKR6|/1-999   200 LQLINDNTATALSYGVFRRKDIN--STAQNVMFYDMGSGSTVCTIVTYQTVKTKEA-GMQPQLQIRGVGFDRTLGGLEMELRLRE 281
sp|Q60432|/1-999   200 LQLINDNTATALSYGVFRRKDIN--STAQNVMFYDMGSGTTVCTIVTYQTVKTKEA-GMQPQLQIRGVGFDRTLGGLEMELRLRD 281
tr|O46067|/1-923   187 LQLINDYAAVALNYGVFHRGEIN--ETAQYFLFYDMGAYKTSAAVVSYQLVKDKQTREINPVVQVLGVGYDRTLGGLEIQLRLRD 269
sp|P36016|/1-881   205 TYLVSEGMSVAVNFVLKQRQFPP--GEQQHYIVYDMGSGSIKASMFSIL--QPEDT-TQPVTIEFEGYGYNPHLGGAKFTMDIGS 284
sp|Q7ZUW2|/1-980   192 LQLINDNTAVALNYGVFRRKDIN--STAQNIMFYDMGSGSTTATIVTYQTVKTKES-GTQPQLQIRGVGFDRTLGGFEMELRLRD 273
tr|C0H5H0|/1-932   227 LGIINGVTAAAIHN-V--HDIPL--NTTKLTMYLDIGSKNINVGIATISFVEKD-K-VRSRSVQVYACESLENNSGNKIDMLLAE 304
sp|Q92598|/1-858   168 LRLMNDMTAVALNYGIYKQDLPSLDEKPRIVVFVDMGHSAFQVSACAFN----------KGKLKVLGTAFDPFLGGKNFDEKLVE 242
sp|Q61699|/1-858   168 LRLMNDMTAVALNYGIYKQDLPNAEEKPRVVVFVDMGHSSFQVSACAFN----------KGKLKVLGTAFDPFLGGKNFDEKLVE 242
sp|P32589|/1-693   169 VRIVNDVTAAGVSYGIFKTDLPEGEEKPRIVAFVDIGHSSYTCSIMAFK----------KGQLKVLGTACDKHFGGRDFDLAITE 243

sp|Q9Y4L1|/1-999   282 RLAGLFNEQRKGQRAKDVRENPRAMAKLLREANRLKTVLSANA-DHMAQIEGLMDDVDFKAKVTRVEFEELCADLFERVPGPVQQ 365
sp|Q9JKR6|/1-999   282 HLAKLFNEQRKGQKAKDVRENPRAMAKLLREANRLKTVLSANA-DHMAQIEGLMDDVDFKAKVTRVEFEELCADLFDRVPGPVQQ 365
sp|Q60432|/1-999   282 HLAKLFNEQRRGQKAKDVRENPRAMAKLLREANRLKTVLSANA-DHMAQIEGLMDDVDFKAKVTRVESEELCADLFERVPGPVQQ 365
tr|O46067|/1-923   270 YLAQEFNALKKT--KTDVTTSPRALAKLFKEAGRLKNVLSANT-EFFAQIENLIEDIDFKLPVTREKLEQLCEDLWPRATKPLEE 351
sp|P36016|/1-881   285 LIENKFLETHPAIRTDELHANPKALAKINQAAEKAKLILSANS-EASINIESLINDIDFRTSITRQEFEEFIADSLLDIVKPIND 368
sp|Q7ZUW2|/1-980   274 HLAKLFNEQKKS--KKDVRDNLRAMAKLLKEAQRLKTVLSANA-EHTAQIEGLMDDIDFKAKVTRSEFEALCEDLFDRVPGPVKQ 355
tr|C0H5H0|/1-932   305 NLRKKFEEKY----NVSIENDKKAMRKLIVAANKAKLLLSAKK-SADVFIESLYNNKSLNESVSRQDFEELIQEVIENMKIPINK 384
sp|Q92598|/1-858   243 HFCAEFKTKY----KLDAKSKIRALLRLYQECEKLKKLMSSNSTDLPLNIECFMNDKDVSGKMNRSQFEELCAELLQKIEVPLYS 323
sp|Q61699|/1-858   243 HFCAEFKTKY----KLDAKSKIRALLRLHQECEKLKKLMSSNSTDLPLNIECFMNDKDVSGKMNRSQFEELCAELLQKIEVPLHS 323
sp|P32589|/1-693   244 HFADEFKTKY----KIDIRENPKAYNRILTAAEKLKKVLSANT-NAPFSVESVMNDVDVSSQLSREELEELVKPLLERVTEPVTK 323

sp|Q9Y4L1|/1-999   366 ALQS----AEMSLDEIEQVILVGGATRVPRVQEVLLKAVGKEELGKNINADEAAAMGAVYQAAALSKAFKVKPFVVRDAVVYPIL 446
sp|Q9JKR6|/1-999   366 ALQS----AEMSLDQIEQVILVGGATRVPKVQEVLLKAVGKEELGKNINADEAAAMGAVYQAAALSKAFKVKPFVVRDAVIYPIL 446
sp|Q60432|/1-999   366 ALQS----AEMSLDEIEQVILVGGATRVPKVQEVLLKAVGKEELGKNINADEAAAMGAVYQAAALSKAFKVKPFVVRDAVIYPIL 446
tr|O46067|/1-923   352 ALAS----SHLSLDVINQVILFGGGTRVPRVQETIKAV-IKQELGKNLNADESATMGAVYKAADLSAGFKVKKFVVKDATLFPLQ 431
sp|P36016|/1-881   369 AVTKQFGGYGTNLPEINGVILAGGSSRIPIVQDQLIKLVSEEKVLRNVNADESAVNGVVMRGIKLSNSFKTKPLNVVDRSVNTYS 453
sp|Q7ZUW2|/1-980   356 ALAA----AEMSMDEIEQVILVGGATRVPKVQDVLLKSVGKEELSKNINADEAAAMGAVYQAAALSKAFKVKPFLVRDAAVFPIQ 436
tr|C0H5H0|/1-932   385 ALEK----GGFQLKDIEALELIGSGWRVPKILNEVTEFFNPLKVGMHLNSDEAVTMGSLYIAAYNSANFRLKDLDYKDIVSNEYH 465
sp|Q92598|/1-858   324 LLEQ----THLKVEDVSAVEIVGGATRIPAVKERIAKFFGK-DISTTLNADEAVARGCALQCAILSPAFKVREFSVTDAVPFPIS 403
sp|Q61699|/1-858   324 LMAQ----TQLKAEDVSAIEIVGGATRIPAVKERIAKFFGK-DVSTTLNADEAVARGCALQCAILSPAFKVREFSVTDAVPFPIS 403
sp|P32589|/1-693   324 ALAQ----AKLSAEEVDFVEIIGGTTRIPTLKQSISEAFGK-PLSTTLNQDEAIAKGAAFICAIHSPTLRVRPFKFEDIHPYSVS 403

sp|Q9Y4L1|/1-999   447 VEFTREVEEEPGIHSLK-HNKRVLFSRMGPYPQRKVITFNRY-SHDFNFHINYGDLGFLGPEDLRVFGSQNLTTVKLKGVGDSF- 528
sp|Q9JKR6|/1-999   447 VEFTREVEEEPGLRSLK-HNKRVLFSRMGPYPQRKVITFNRY-SHDFNFHINYGDLGFLGPEDLRVFGSQNLTTVKLKGVGESF- 528
sp|Q60432|/1-999   447 VEFTREVEEEPGVRSLK-HNKRVLFSRMGPYPQGKVITFNRY-SHDFNFHINYGDLGFLGPEDLRVFGSQNLTTVKLKGVGESL- 528
tr|O46067|/1-923   432 VSFERDPGDGAAVK----QVKRALFALMNPYPQKKVITFNKH-TDDFEFYVNYADLDRYSKEEIAALGSLNVTKVQLKQVKELL- 510
sp|P36016|/1-881   454 FKLSNESELY------------DVFTRGSAYPNKTSILTNTTDSIPNNFTIDLFENGKL-------FET---ITVNSGAIKNSY- 515
sp|Q7ZUW2|/1-980   437 VEFSRETEEEDGVKSLK-HNKRILFQRMAPYPQRKVITFNRY-IDDFVFYINYGDLSFLSEQDMKVFGSQNLTTVKLSGVGSSF- 518
tr|C0H5H0|/1-932   466 ILVNTDEEENNTTNEEKVNIKKELVNYNSRYPHNKNVILTYK--DNLKFSVY---------ENGKIINEYVLGNL-----DNAIK 534
sp|Q92598|/1-858   404 LIWNHDSEDTEGVH--------EVFSRNHAAPFSKVLTFLRR--GPFELEAFYSDPQGVPYPE-AKIGRFVVQNVSA-------- 469
sp|Q61699|/1-858   404 LVWNHDSEETEGVH--------EVFSRNHAAPFSKVLTFLRR--GPFELEAFYSDPQGVPYPE-AKIGRFVVQNVSA-------- 469
sp|P32589|/1-693   404 YSWDKQVEDEDHM---------EVFPAGSSFPSTKLITLNRT--GDFSMAASYTDITQLPPNTPEQIANWEITGVQL-------- 469

sp|Q9Y4L1|/1-999   529 KKY--PDYESKGIKA--HFNLDESGVLSLDRVESVFETLV-EDSAEEESTLTKLGNTISSLF-GGGTTPDAKENGTDTVQEEEES 607
sp|Q9JKR6|/1-999   529 KKY--PDYESKGIKA--HFNLDESGVLSLDRVESVFETLV-EDSPEEESTLTKLGNTISSLF-GGGTSSDAKENGTDAVQEEEES 607
sp|Q60432|/1-999   529 KKY--PDYESKGIKA--HFNLDESGVLSLDRVESVFETLV-EDSPEEESTLTKLGNTISSLF-GGGTSSDAKENGTDAVQEEEES 607
tr|O46067|/1-923   511 EKSKKELVDNKGIKA--YFYLDDSGIFRCTGVEYVYEKQKPEDDADEDSTLSKFGSTLSKLF-TKEGE-EKK------------- 578
sp|P36016|/1-881   516 --S--SDKCSSGVAYNITFDLSSDRLFSIQEVNCICQSEN-D--IGNSKQIKNKGSRLA--F----------------------- 568
sp|Q7ZUW2|/1-980   519 KKH--SDAESKGIKA--HFNMDESGVLILDRVESVFETIV-EEK-EEESTLTKLGNTISSLF-GGGSS-EPSANVTEPVTDEEEV 595
tr|C0H5H0|/1-932   535 SKY--EHLGTPKLNL--KFHLDKFGILSLDKVLVVYEEQK-D-----G----------------AGDTKDNKKEGDE-------- 585
sp|Q92598|/1-858   470 --Q--KDGEKSRVKV--KVRVNTHGIFTISTASMVEKVPT-E-----ENEMSS-EADMECLNQRPPENPDTDKN----VQQD--- 534
sp|Q61699|/1-858   470 --Q--KDGEKSRVKV--KVRVNTHGIFTISTASMVEKVPT-E-----EEDGSSLEADMECPNQRPTESSDVDKN----IQQD--- 535
sp|P32589|/1-693   470 --P--EGQDSVPVKL--KLRCDPSGLHTIEEAYTIEDIEV-E-----EP------------------------------------ 506

sp|Q9Y4L1|/1-999   608 PAEGSK--DEPGEQVELKEEAEAPVEDGS--QP-PPPEPKG--DAT-PEGEKATEKENGDKS---EAQKPSEKAEAGPEGVAPAP 681
sp|Q9JKR6|/1-999   608 PAEGSK--DEPAEQGELKEEAEPPAEETS--QP-PPSEPKG--DAA-REGEKPDEKESGDKP---EAQKPNEKGQAGPEGAAPAP 681
sp|Q60432|/1-999   608 PTEGSK--DEPGEQGDLKEETEAPVEDTS--QP-PPPEPKG--DAA-PEGEKPDEKESGGKS---EAQKPEEKGQSGPEGVPPAP 681
tr|O46067|/1-923   579 ---------DNSEQEE----------------------------AA-NAGEEPSKSEDNEKAKEEDASK-EQ-----------KS 613
sp|P36016|/1-881       -------------------------------------------------------------------------------------
sp|Q7ZUW2|/1-980   596 TPEAGKEQDQPEKQEETVQEK-PETEEGKEAEP-QAEEQKE--DKEKAENQGETESEKTEKP---EEKTTDE-----------EK 662
tr|C0H5H0|/1-932   586 --ENNN--NNNNEEI------NKDDDTNN--NKSDDEQNKGDENKS-NDENKE-------------------------------N 626
sp|Q92598|/1-858   535 ---N----SEAGTQPQV----QTDAQQTS--QSPPSPELTSEENKI-PDADKA-------------------------------N 574
sp|Q61699|/1-858   536 ---N----SEAGTQPQV----QTDGQQTS--QSPPSPELTSEESKT-PDADKA-------------------------------N 575
sp|P32589|/1-693   507 ---------------------------------IPLPED-----AP-EDAE---------------------------------Q 519
```

```
sp|Q9Y4L1|/1-999  682 EGEKKQKPARK-RRMV-EEIGV------ELVV--LD--LPDLPE--DKLAQSVQKLQDLTLRDLEKQEREKAANSLEAFIFETQD 752
sp|Q9JKR6|/1-999  682 EEDKKPKPARK-QKMV-EEIGV------ELAV--LD--LPDLPE--DELARSVQKLEELTLRDLEKQEREKAANSLEAFIFETQD 752
sp|Q60432|/1-999  682 EEEKKQKPARK-QKMV-EEIGV------ELAV--LD--LPDLPE--DELARSVKKLEDLTLRDLEKQEREKAANSLEAFIFETQD 752
tr|O46067|/1-923  614 EESTKQDTEAK-NETI-KLVTVKSPVTYESQT--QF--VVPLVG--SAYDQSVAKLAAINKAEEQRVRLESAFNALEAHIIEVQQ 690
sp|P36016|/1-881  569 ------------------------------TS--EDVEIKRLSP--SERSRLHEHIKLLDKQDKERFQFQENLNVLESNLYDARN 619
sp|Q7ZUW2|/1-980  663 EADMKPKLQKK-SKIS-ADIAV------ELEV--ND--VLDPSA--EDMEGSKKKLQDLTDRDLEKQEREKTLNSLEAFIFETQD 733
tr|C0H5H0|/1-932  627 EE-NKQNGEKKKNDIIKHNIPI------EFQT--RN--IKPLPLTFEEIKEKKEILKNLDEHDIDIFLKSEKKNTLESFIYETRS 700
sp|Q92598|/1-858  575 EKKVDQPPEAK-KPKI-KVVNV------ELPIEANL--VWQLGK--DLLNMYIETEGKMIMQDKLEKERNDAKNAVEEYVYEFRD 647
sp|Q61699|/1-858  576 EKKVDQPPEAK-KPKI-KVVNV------ELPVEANL--VWQLGR--DLLNMYIETEGKMIMQDKLEKERNDAKNAVEECVYEFRD 648
sp|P32589|/1-693  520 EF-KKVTKTVK-K--------D------DLTIVAH---TFGLDA--KKLNELIEKENEMLAQDKLVAETEDRKNTLEEYIYTLRG 583

sp|Q9Y4L1|/1-999  753 KLYQPEYQEVSTEEQREEISGKLSAASTWLEDEGVGATTVMLKEKLAELRKLCQGLFFRVEERKKWPE-RLSALDNLLNHSSMFL 836
sp|Q9JKR6|/1-999  753 KLYQPEYQEVSTEEQREEISGKLSATSTWLEDEGFGATTVMLKDKLAELRKLCQGLFFRVEERRKWPE-RLSALDNLLNHSSIFL 836
sp|Q60432|/1-999  753 KLYQPEYQEVSTEEQREEISGKLSATSTWLEDEGFGATTVMLKEKLAELKKLCQGLFFRVEERRKWPE-RLSALDNLLNHSSIFL 836
tr|O46067|/1-923  691 KLDEESYAKCATAEEKEKLLAECSTLGEWLYEDLEDPKAEIYEEKLAQLKKLSNVFLARHWEHEERPE-AIKALKGMIDGAEKFL 774
sp|P36016|/1-881  620 LLMDDEVMQNGPKSQVEELSEMVKVYLDWLEDASFDTDPEDIVSRIREIGILKKKIELYMDS-AKEPL-NSQQFKGMLEEGHKLL 702
sp|Q7ZUW2|/1-980  734 KLYQDEYQAVVTEEEKEQISGRLSVASSWMDEEGYRAGTKLLKEKLSELKKLCKGMFFRVEERKKWPD-RLAALDSMLNHSNIFL 817
tr|C0H5H0|/1-932  701 KMKQDIYKQVTKEETRNEYLNKLEEYEDWLYTEKDEP-LENVSNKIHELQDIYNPIKERAEELQVRDK-IIEETNKKIQEMIE-- 781
sp|Q92598|/1-858  648 KLCG-PYEKFICEQDHQNFLRLLTETEDWLYEEGEDQAKQAYVDKLEELMKIGTPVKVRFQEAEERPK-MFEELGQRLQHYAKIA 730
sp|Q61699|/1-858  649 KLCG-PYEKFICEQEHEKFLRLLTETEDWLYEEGEDQAKQAYIDKLEELMKMGTPVKVRFQEAEERPK-VLEELGQRLQHYAKIA 731
sp|P32589|/1-693  584 KLEE-EYAPFASDAEKTKLQGMLNKAEEWLYDEGFDSIKAKYIAKYEELASLGNIIRGRYLAKEEEKKQAIR------------- 654

sp|Q9Y4L1|/1-999  837 KGARLIPE----MDQIFTEVEM---TTLEKVINE-------------TWAWKN------ATLAEQAKL-PAT------------- 881
sp|Q9JKR6|/1-999  837 KGARLIPE----MDQVFTEVEM---TTLEKVIND-------------TWAWKN------ATLAEQAKL-PAT------------- 881
sp|Q60432|/1-999  837 KGARLIPE----MDQIFTEVEM---TTLEKVIND-------------TWAWKN------ATLAEQAKL-PAT------------- 881
tr|O46067|/1-923  775 VTGRNLTKDTNPEKDVFTQVEI---DTLDKVITE-------------TNAWLK------TETAAQKKL-AKN------------- 823
sp|P36016|/1-881  703 QAIETHKNT---VEEFLSQFETEFADTIDNVREEFKKIKQPAYVSKALSTWEETLTSFKNSISEIEKFLAKNLFGEDLREHLFEI 784
sp|Q7ZUW2|/1-980  818 KSARLIPE----SDQIFTDVEL---KTLEKVINE-------------TITWKN------ETVAEQEKL-SPT------------- 862
tr|C0H5H0|/1-932  782 -KIKDLSE----KKPWA-AETI---KMVKDSLDK-------------EVQWWN------HAQEEQKKL-DNY------------- 824
sp|Q92598|/1-858  731 ADFRNKDE----KYNHIDESEM---KKVEKSVNE-------------VMEWMN------NVMNAQAKK-SLD------------- 775
sp|Q61699|/1-858  732 ADFRGKDE----KYNHIDESEM---KKVEKSVNE-------------VMEWMN------NVMNAQAKR-SLD------------- 776
sp|P32589|/1-693      -------------------------------------------------------------------------------------

sp|Q9Y4L1|/1-999  882 ------------EKPVLLSKDIEAKMMALDR--EVQYLLNK---AKFTKPRP---RPKDKNGTRAEPPLNASASDQGEKVIPPAG 946
sp|Q9JKR6|/1-999  882 ------------EKPVLLSKDIEAKMMALDR--EVQYLLNK---AKFTKPRP---RPKDKNGTRAEPPLNASAGDQEEKVIPPAG 946
sp|Q60432|/1-999  882 ------------EKPVLLSKDIEAKMMALDR--EVQYLLNK---AKFTKPRP---RPKDKNGTRTEPPLNATAGDQEEKVIPPAG 946
tr|O46067|/1-923  824 ------------ADIRLTVKDITDKMSLLDR--EVKYLVNK---IKIWKPKV---KPAAEKEKKKEEEVVASGSGDDTK------ 882
sp|P36016|/1-881  785 KLQFDMYRTKLEEKLRLIKSGDESRLNEIKKLHLRNFRLQKRKEEKLKRKLE---QEKSRNNNETESTVINSA-DDKTTIVND-- 863
sp|Q7ZUW2|/1-980  863 ------------VKPVLLSKDIEAKLSLLDR--EVNYLLNK---AKFAKPKP---KDKAKDKNSTSESSKANSTDDAEKVIPPKT 927
tr|C0H5H0|/1-932  825 ------------TAPFFKHKDVQLKFKSIQM--LIKTLD------KLKKPVE---KKEDKKNTDNQNE-NTSKQDAGADKN---- 881
sp|Q92598|/1-858  776 ------------QDPVVRAQEIKTKIKELNN--TCEPVVT--------QPKP---KIESPKLERTPNGPNI---DKKE------- 825
sp|Q61699|/1-858  777 ------------QDPVVRTHEIRAKVKELNN--VCEPVVT--------QPKP---KIESPKLERTPNGPNI---DKK-------- 825
sp|P32589|/1-693  655 ------------------SKQEASQMAAMAE-----KLAA--------QRKAEAEKKEEKK------------------------ 684

sp|Q9Y4L1|/1-999  947 Q--TEDAEPISEPEKVETGSEP-GDTEPLELGGPGAEPEQKEQ-STG-QKRPLKNDEL-                           999
sp|Q9JKR6|/1-999  947 Q--TEEAKPILEPDKEETGTEP-ADSEPLELGGPGAGPEQEEQ-SAG-QKRPSKNDEL-                           999
sp|Q60432|/1-999  947 Q--PEEAKPILEPDKEETTTEP-TDSEPLELGGPGAESEPKEQ-TAG-QKRSSKNDEL-                           999
tr|O46067|/1-923  883 ---SEDAEQQQEQA---TKEEQ-QEQEPVDEITPTP---------AE-EETKTPHSEL-                           923
sp|P36016|/1-881  864 K--TT------------------------------------ES-NPS-SEEDILHDEL-                           881
sp|Q7ZUW2|/1-980  928 EDGAEKVKPAEEPPVVE---EK-AEETILELN-PAENTDDKTESTESSKSENHIEDEL-                           980
tr|C0H5H0|/1-932  882 ---HNTTENQNEQSAQNQNNENNDDNQNNEHD--ANQSSNDEQ-NKN-DGASDQKDE-L                           932
sp|Q92598|/1-858  826 ----EDLEDKNN-----FGAEP---------------PHQNGE-CYP-NEKNSVNMDLD                           858
sp|Q61699|/1-858  826 ----EDLEGKNN-----LGAEA---------------PHQNGE-CHP-NEKGSVNMDLD                           858
sp|P32589|/1-693  685 ----------------------------------------------D-TE-GDV--DMD                           693
```

Q9Y4L1 - Human GRP170
Q9JKR6 - Mouse GRP170
Q60432 – hamster GRP170
O46067 – Drosophila GRP170
P36016 – yeast GRP170
Q7ZUW2 – Zebra fish GRP170
C0H5H0 – Plasmodium GRP170
Q92598 – human Hsp110
Q61699 – mouse Hsp110
P32589 – Yeast Sse1

| |
|---|
| > 80 % |
| > 60 % |
| > 40% |
| < 40% |

Colors indicate the percentage of residues that agree with the consensus sequence for that column.
